# Spontaneous back-pain alters randomness in functional connections in large scale brain networks

**DOI:** 10.1101/596163

**Authors:** Gurpreet S. Matharoo, Javeria A. Hashmi

## Abstract

We use randomness as a measure to assess the impact of evoked pain on brain networks. Randomness is defined here as the intrinsic correlations that exist between different brain regions when the brain is in a task-free state. We use fMRI data of three brain states in a set of back pain patients monitored over a period of 6 months. We find that randomness in the task-free state closely follows the predictions of Gaussian orthogonal ensemble of random matrices. However, the randomness decreases when the brain is engaged in attending to painful inputs in patients suffering with early stages of back pain. A persistence of this pattern is observed in the patients that develop chronic back pain, while the patients who recover from pain after 6 months, the randomness reverts back to a normal level.

**Author Summary:** Back-pain is a salient percept known to affect brain regions. We studied random correlations in brain networks using random matrix theory. The brain networks were generated by fMRI scans obtained from a longitudinal back-pain study. Without modelling the neuronal interactions, we studied universal and subject-independent properties of brain networks in resting state and two distinct task states. Specifically, we hypothesized that relative to the resting state, random correlations would decrease when the brain is engaged in a task and found that the random correlations showed a maximum decrease when the brain is engaged in detecting back pain than performing a visual task.

## Introduction

Chronic pain represents a major clinical, social, and economic problem for societies worldwide. The principal complaint is of unremitting physical pain that does not abate with standard analgesics(1–3). The experience of pain is quite different across the population and persists for different durations between individuals. Pain is in essence a threat signal that we localize to a part of the body in the form of an unpleasant sensation. This sensation accompanies a strong negative emotion that works as an aversive signal which is necessary for learning proper avoidance behaviors. In some people, this signal becomes accentuated and tends to persist for long periods of times extending over months to years. These individuals very often show no signs of tissue damage or underlying pathology in the site where they are feeling pain. Brain imaging studies suggest that chronic pain alters the nervous system so that the brain perceives persistent pain due to maladaptive processes in the brain. An expedient approach for understanding these maladaptive processes is to observe how back pain transitions to a chronic form.

Thus, we know that in some patients, persistent back pain is acute and persists for a few weeks to be classified as subacute back pain (or SBP). This early stage of persistent back pain remits in some individuals, while for others, it persists for months to years and this enduring back pain is classified as chronic (Chronic Back Pain or CBP). The reasons and neural mechanisms due to which back pain transitions from subacute to chronic is still ambiguous, and the pursuit to find neurological reasons for this transition is central to contemporary pain research. In recent years, there have been successful attempts in relating CBP to specific brain activity(4) whereby neuroimaging method of functional Magnetic Resonance Imaging (fMRI) is used to study the correlations between CBP and brain activity. More recently, it has also been shown that chronification of back pain shifts the brain activity from nociceptive to emotional circuits, thereby impacting patients with physiological disorders such as depression and impacting their overall quality of everyday life(3).

fMRI makes use of the fact that neuronal activity is partly coupled with increases in blood flow in the observed parts of the brain and it images these changes as a haemodynamic response to brain activity. This particular form of fMRI is also referred to as blood-oxygenation-level-dependent (BOLD) fMRI and it offers high spatial resolution. A useful adaptation of this approach is to measure how slow temporal fluctuations (0.01-0.15 HZ) are between different brain regions and this statistical dependency is referred to, more generally, as functional connectivity. The network properties that emerge from large-scale correlations has been shown to be altered in neuropsychiatric and chronic conditions such as CBP(4–9). It is still a challenge to understand the dynamic transition of brain between different states as a result of back-pain. It is because brain is a fairly complex system whereby neurons are constantly interacting with each other often resulting in higher brain functions(10,11) and in the formation of functional networks, even in the absence of any stimuli. Though large-scale functional connectivity is often studied using clustering techniques or principles of graph theory(12), there is a need to apply the concepts and methodologies developed in the context of the theory of random matrices for observing systematic transitions in brain states.

Random Matrix Theory (RMT) was originally developed in the nuclear physics applications, where nuclei can have many possible states and energy levels and, and their interactions are too complex to be described accurately. In such a scenario, one settles for a model that captures the statistical properties of the energy spectrum. RMT finds extensive applications in the statistical studies of various complex systems such as quantum chaotic systems, complex nuclei, atoms, molecules, disordered mesoscopic systems(13–21), atmosphere(22), financial applications(23), network forming systems(24,25), amorphous clusters(26–29), biological networks(30,31), etc. In recent years, RMT has also been applied towards brain network studies in studying universal behavior of brain functional connectivity and has been effective in detecting the differences in resting state and visual stimulation state(32,33). Recently, attempts using RMT have also been made in brain functional network studies on attention deficit hyperactivity disorder (ADHD)(34). RMT makes use of the fact that true information of the system is contained in the eigenvalues of a correlation matrix. Specifically, for brain networks, the eigenvalues represent the level of functional connectivity between different regions of interest (ROIs) in brain, and larger eigenvalues contain information about significant correlations (or strong connectivity), and therefore, about processes in brain. Recent studies have shown that ROIs in brain are correlated. Furthermore, these correlations closely follow the predictions of Gaussian Orthogonal Ensemble (GOE) of random matrices when the brain is in a state of rest (fully-conscious). The clearest indication so far has come from EEG data(32), which further attributes the observed deviation from GOE predictions to visual stimulation; that is, true information. Other recent studies(33,34) also point to similar information, however, the overall findings are unclear. We hereby propose a hypothesis where, we refer to these observed correlations as random correlations, or in general, randomness, that exists at any given instant in brain network. When the brain is engaged in a task, this randomness would be expected to decrease, as brain regions would be connected in a coherent fashion relative to a task-free or resting state. These random correlations reach their normal levels at resting state. Thus, RMT may offer a principled approach for measuring systematic changes in randomness that occur in brain networks during perception and cognition.

Here we investigate whether the brain demonstrates a greater deviation from GOE predictions when it is engaged in detecting threats or experiencing discomfort from pain relative to perception of innocuous stimuli. Since the ability to properly detect and perceive pain is fundamental for survival, attending to pain can be expected to add systematic changes in brain connectivity and thus reduce random correlations in brain networks. On the other hand, maladaptive processing of pain inputs during a chronic stage of back pain may show a different behavior, relative to the SBP state. The ability to distinguish these two states using an integrative approach such as RMT could be useful for improving chronic pain diagnosis and prognosis and also for understanding the abnormalities in brain properties that contribute to CBP.

## Materials and methods

### Dataset and Tasks

We use fMRI data available on the open access data sharing platform for brain imaging studies of human pain (www.openpain.org). The complete dataset is a part of 5-year longitudinal study of transition to chronic back pain in which 120 patients were recruited initially. All the participants were trained to perform two tasks using finger-span device with which they provided continuous pain ratings(3,4). This device consisted of a potentiometer in which voltage was digitized. During the brain imaging sessions, the device was synchronized and time-stamped with fMRI image acquisition and connected to a computer providing visual feedback of the pain ratings(35). We use data acquired from three different states, a) A state of rest in which the participants are not thinking about any one thing in particular (RS); b) A state of focusing and rating spontaneous changes in back pain (SP); and, c) A control state in which they are rating changes in length of a visual bar (SV).

### MRI data acquisition

The data for all participants and visits was collected by a 3T Siemens scanner. At first, MPRAGE type T_1_ anatomical brain images were acquired followed by fMRI scans on the same day with the following parameter details given in *Hashmi et al*(3):

EPI sequence: voxel size 1 × 1 ×1 MM, Repetition time=2500MS; Echo Time=3.36MS; Flip angle = 9degrees; In-Plane matrix resolution 256 × 256; slices 160, filed of view, 256mm. Functional MRI scans were acquired on the same day as the T1 scan and MPQVAS measures: multi-slice T2*-weighted EPI images with repetition time=2.5s, echo time=30ms, flip angle =90 degree, number of volumes =244, slice thickness =3mm, in-plane resolution =64 × 64.

### Pre-processing of fMRI data

We use Freesurfer, FMRIB Software Library (FSL) v5.0, and Analysis of Functional Neuro-Images (AFNI) software to preprocess the data similar to procedures adapted for the 1000 Functional Connectomes project(36). Data were slice time corrected, motion corrected, temporally band-pass filtered, and then further filtered to remove linear and quadratic trends using AFNI. Complete details of the preprocessing procedure are given in(37). The registration was performed using FMRIB’s Linear and non LINEAR Image Registration Tools for transformations from native functional and structural space to the Montreal Neurological Institute MNI152 template with 2 × 2 × 2 resolution, with further details given in(37).

### Anatomical parcellation and construction of correlation matrix

The brain is anatomically parcellated by *an optimization of the Harvard/Oxford parcellation scheme* (OHOPS)(38). In this scheme, the anatomical partitioning of cingulate, medial and lateral prefrontal cortices of Harvard Oxford Atlas was increased and in addition, anatomical partitioning of insular label was also performed, and the single Region of Interest (ROI) spanning the entire insula in Harvard Oxford Atlas was further subdivided based on a previous scheme(39). The complete OHOPS consisted of a total of 131 regions(38). Each ROI was designated as a node and the BOLD time series were extracted from each node and averaged to generate 131 time series for each subject. Following this, the whole brain networks were constructed, and network measures were assessed using the Brain Connectivity Toolbox, with formulae used for calculating network measures described in(40). The brain networks are usually assortative in nature(41,42).

For each patient, the BOLD time series in each region was correlated with every other region to create a 131 × 131 symmetric correlation matrix based on Pearson’s correlation coefficients given by:

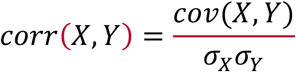

or, which can be re-written as:

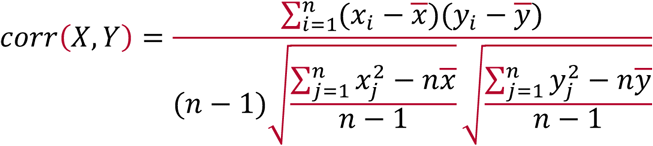

Such correlation matrices are not only symmetric, but they are also positive semi-definite(43), with all eigenvalues being non-negative. This correlation matrix is then diagonalized and eigenvalues (λ) are obtained. In the present case, few eigenvalues are zeros, and remaining have positive values. It must be remembered that not all ROIs are a part of active brain network at a given time and hence, very small eigenvalues are usually ignored, and the related correlations are unimportant from functional connectivity perspective.

### Unfolding of data

Fluctuations around the eigenvalue spectra are studied using standard methods of RMT. The first step is to unfold the data, meaning, the eigenvalues are arranged in an increasing (cumulative) order and are then mapped using an analytical function in such a way that the average spacing between two successive eigenvalues is unity. This ensures all the eigenvalues are on same-footing. The analytical fitting function used for unfolding need not be unique and, is generally different for different systems(25–29). For this study, the eigenvalue spectra of all the correlation matrices generated is approximated extremely well by a function of the form

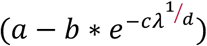

where a, b, c, and d, are best-fit parameters and λ is the eigenvalue. Figure 1 shows a plot of the cumulative eigenvalue density along with the analytical fitting function. We leave out a small portion of eigenvalues at both ends in order to achieve the best fit, something which has been a standard practice in other works(25–29). We deal with unfolded eigenvalues from this point onwards.

**Fig. 1:**
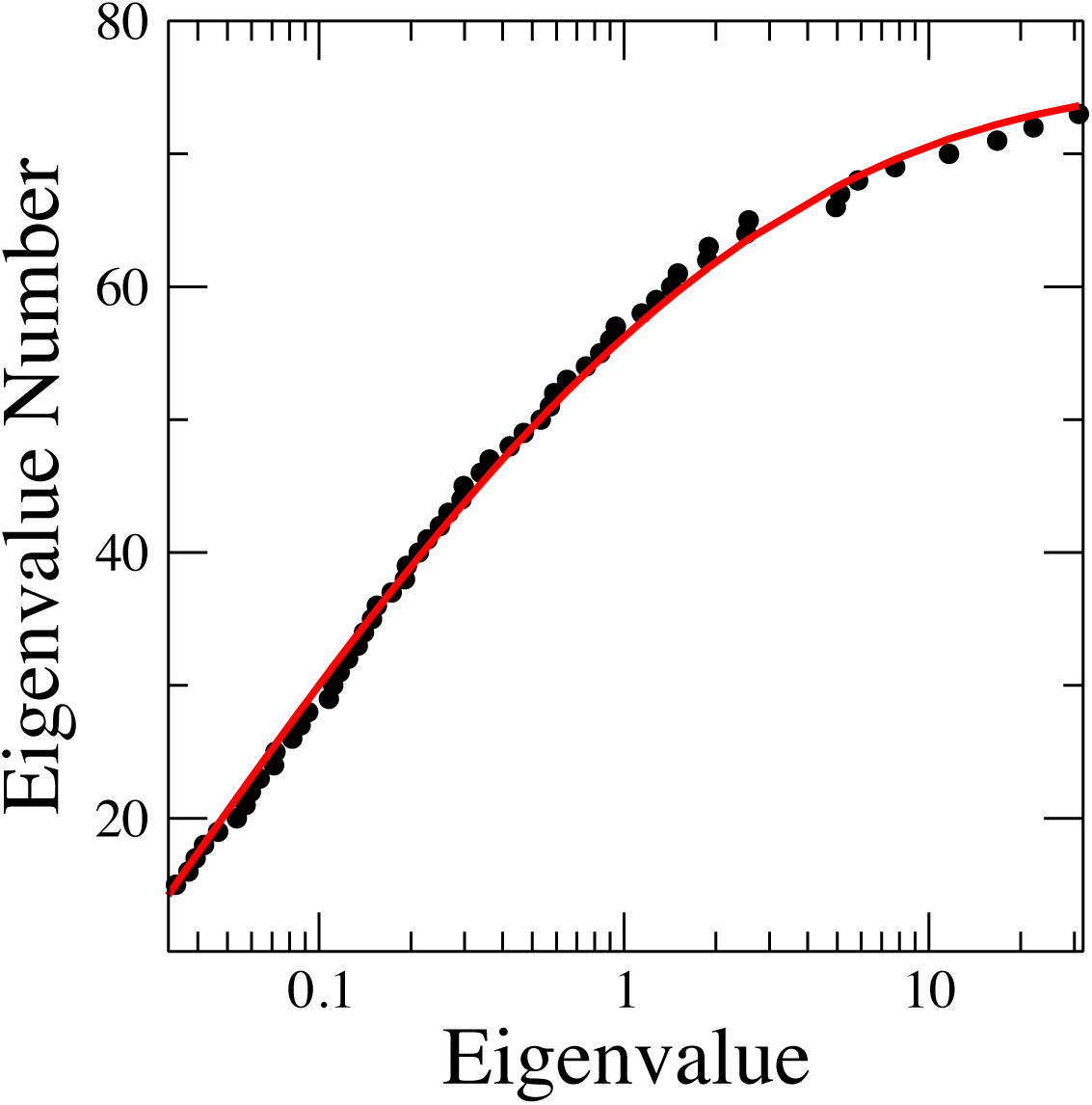
Eigenvalue number vs eigenvalue (λ) for a typical spectrum. Filled circles (black): Data. Continuous line (red): The best-fit using the functional form described in text.

## Results

We report the spectral statistics fluctuation properties of the eigenvalue spectra in the three brain states in individuals who were suffering with SBP (back pain for < 3 months). We also track what these properties looked like after 6 months in the group of individuals with SBP with persisting back pain(3,4,7,44). Patients had all been pain free for one year prior to their subacute pain episode and had no history of any mental illness including depression. The individual details of patients are also available online on the data sharing platform. It must also be stated that none of the data from available subjects was excluded from the analysis.

### Visit 1

For visit 1, 68 SP and SV scans are available. In addition, there are 27 RS scans available for visit 1. Analysis of randomly picked individual eigenvalue spectra indicate that brain-states have fluctuation properties associated with the Gaussian orthogonal ensemble (GOE) of random matrices. To improve statistics, we combine information from all unfolded data. Figure 2a shows the normalized nearest-neighbor spacing distribution (NNSD) [p(s)] for RS, SP, and SV scans for visit 1. Here, *s* is the eigenvalue spacing. Superimposed is the GOE result, which is also approximated by Wigner’s surmise as:

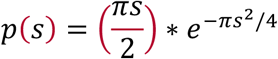

**Fig. 2:**
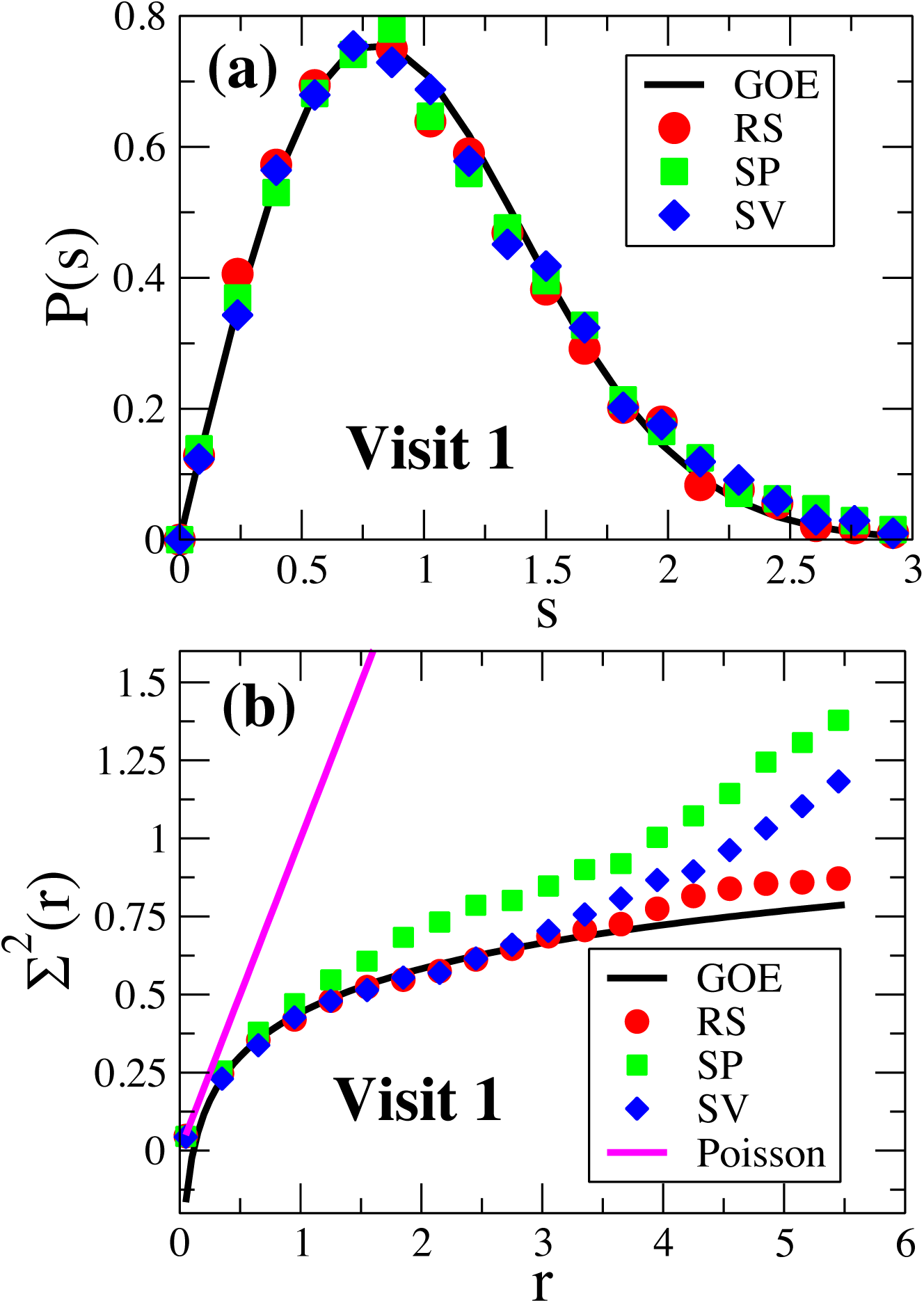
(a) Normalized neighbor spacing *(s)* vs probability density *p(s)* for resting state (red circles), spontaneous pain (green squares), and standard visual (blue diamonds) scans for Visit 1. Solid line is the GOE prediction.; (b) Variance of the number of levels in intervals of length *r* shown as a function of *r* for resting state (red circles), spontaneous pain (green squares), and standard visual (blue diamonds) for Visit 1. Black line represents GOE prediction and magenta line represents Poisson distribution.

For all the cases, we find a good agreement with GOE. A single-valued indicator that follows the p(s) function is the variance of nearest-neighbor spacing. We find this number between 0.297 and 0.320 for all the cases, which is quite close to 0.286, the number for GOE(26–28). This agreement could be explained due to the fact that NNSD captures the correlations that exists between successive eigenvalues and does not have information about the long-range correlations. Short-ranged correlations, especially between the nearest-neighbors are quite strong, and hence not altered substantially by both, visual (SV) and pain-rating (SP) tasks. This result is also consistent to other brain-network studies(32–34,42) and hence, further strengthens the belief that there exists strong, stimuli-resistant random correlations between nearest-neighbors in the brain network.

Next, we take a look at the long-range (or higher order) random correlations. For this, we measure Σ^2^(r), the variance of the number of levels n(r) within an interval of length *r*. The theoretical result for GOE is:

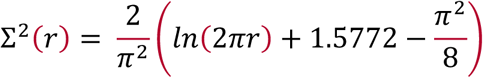

The number variance is quite sensitive to changes, and is extremely sensitive to small systematic errors in the approximation to the analytical function used during unfolding(26,27). Contribution of any such error to Σ^2^(r) grows as *r*^*2*^, whereas the GOE prediction for Σ^2^(r) grows as *ln(r)*(29). In Figure 2b, we plot Σ^2^(r) for RS, SP, and SV scans along with GOE and Poisson [Σ^2^(r) = r] distributions for visit 1. We observe that RS agrees with the GOE prediction over greatest domain, whereas we see deviations for SV and SP scans with SP scans showing maximum deviation. This deviation is attributed to the relative tasks the subjects are performing in each case, with the pain-rating task showing maximum deviation.

### Visit 4

At visit 4, which was approximately 6 months after visit 1, some patients recovered from persistent back-pain as a result of spontaneous remission of the condition (recovering group), others experienced a persistence in their back-pain, and represent the group who have developed CBP (persistent group). To define SBP persistent group, we separate participants with pain persisting for 6 months from those that recovered (SBP recovering) based on self-report of pain ratings observed using McGill Pain Questionnaire Visual Analogue Scale (MPQVAS). We compare the MPQVAS value at visit 1 with visit 4. If the pain rating value of a particular subject decreases by 30% or more, the subject is classified as “Recovering”, else, it is classified as “Persistent”. Based on this classification, we have 18 RS, 17 SP, and 23 SV scans for Persistent group and 18 RS, 19 SP, and 28 SV scans for Recovering group.

Figure 3 shows NNSD for Persistent and Recovering groups. Both the plots show agreement with GOE predictions; an indicator of strong nearest-neighbor random correlations. Figure 4 shows plots of Σ^2^(r) for Persistent and Recovering groups. In both the cases, we find RS scans staying close to GOE predictions. However, we find a striking difference between SP and SV scans in the two cases. For the Persistent group, both SP and SV scans show deviations from the theory, with SP scans showing greater deviations than SV scans. For the Recovering group, both SP and SV scans match GOE predictions over a larger domain, and undistinguishable from RS scans.

**Fig. 3:**
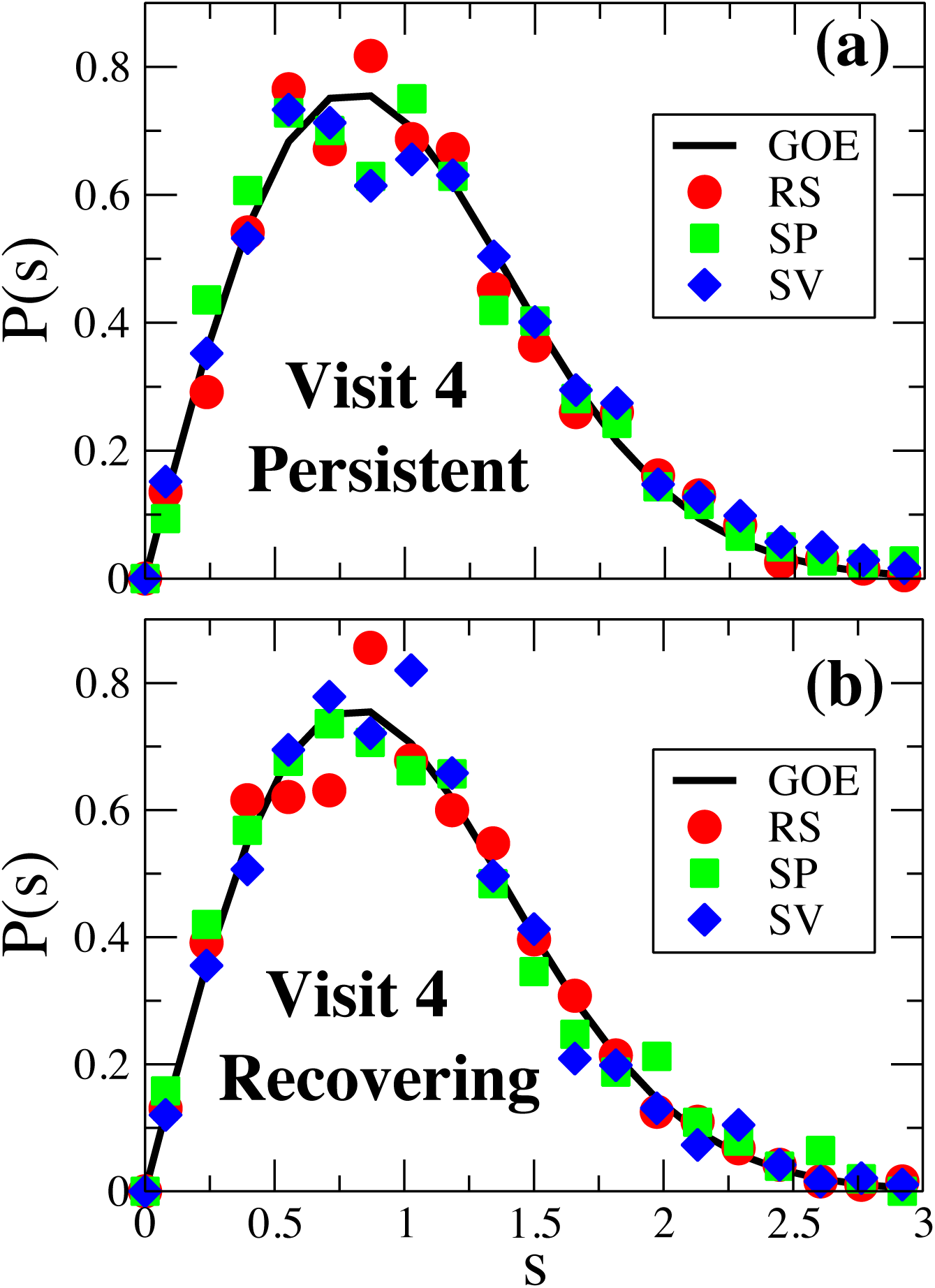
Normalized neighbor spacing *(s)* vs probability density *p(s)* for resting state (red circles), spontaneous pain (green squares), and standard visual (blue diamonds) scans for (a) Persistent, and (b) Recovering groups in visit 4. Solid line is the GOE prediction.

**Fig. 4:**
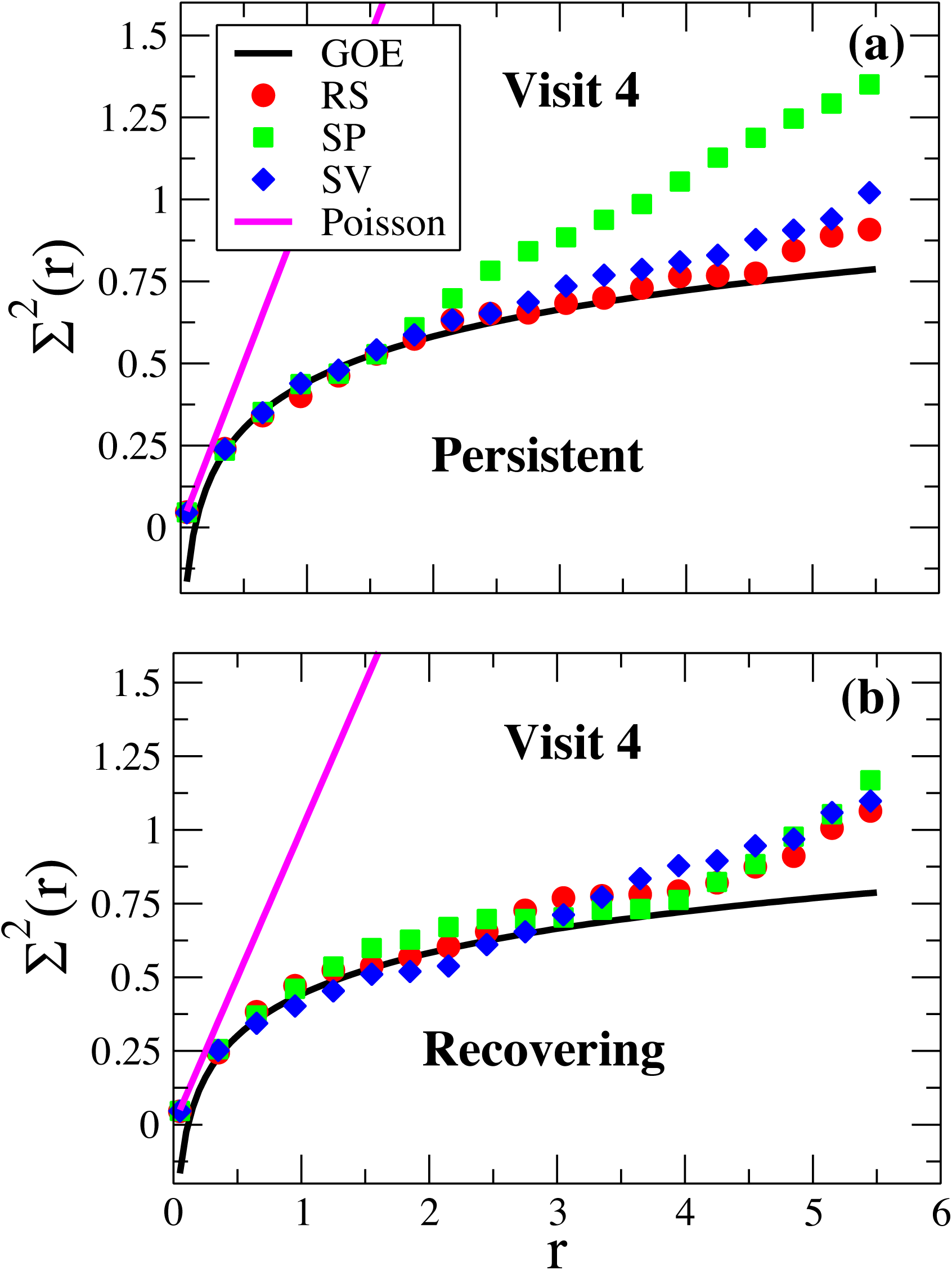
Variance of the number of levels in intervals of length *r* shown as a function of *r* for resting state (red circles), spontaneous pain (green squares), and standard visual (blue diamonds) for (a) Persistent, and (b) Recovering groups in visit 4. For both visits, black line represents GOE prediction and magenta line represents Poisson distribution.

## Discussion

The present study demonstrates that RMT is able to differentiate between two different tasks within the same subject. We find a pattern consistent with our hypothesis, with randomness decreasing when the brain is focused on attending to pain triggered in the back of their body. Here, GOE line represents maximum randomness and Poisson represents no randomness. However, due to the complexity of the experimental design, there could be many possible conjectures (including their combinations) explaining these observations.

First, as the patients are performing a pain-rating task, whereby they are focusing on the back and reporting the ratings, the observed SP deviations could be attributed to back-pain. As it known from earlier studies that salient percepts such as pain are known to require more brain areas to be engaged than visual stimulation, we see an increased deviation for SP scans relative to SV scans in all the cases(45–47). As more brain regions are engaged in attending to pain, hence relative randomness between them decreases. At Visit 1, all patients report back-pain, whereas at Visit 4, only a subset of them report back-pain, and because their MPQVAS ratings demonstrate chronification of pain, the Persistent group continues to experience back-pain over many months. Hence, this continued deviation of SP scans at Visit 4 in the persisting CBP group is a reflection of chronified pain that continues to affect the GOE pattern. Second possible conjecture is the saliency between the tasks themselves. While visual tasks are relatively easy to perform, pain-rating tasks could be much difficult as back-pain events are generally random. Hence, more attention is needed to perform these tasks, and thereby, we observe a decrease in randomness between the brain regions involved in these tasks.

The present study also provides some useful insights on the connectivity states of resting state of brain. Previous spectral studies using random matrix theory on quenched (local minima on potential energy landscape) normal modes of network-forming liquids (Water)(25) and amorphous systems (clusters and periodic systems both two-and three dimensions)(26–29) have demonstrated that the fluctuations around the mean spectral densities follow GOE. For normal modes that are not necessarily quenched, this agreement is not perfect, but gets better with increasing density(24). While it is beyond the present work to prove, and further research is needed along these lines, we propose an ansatz that resting state corresponds to local energy minima whereby the intrinsic correlations obey GOE and conditions like back-pain can be viewed as a perturbation in system dynamics resulting in a shift away from stable local minimum. It also remains an open question whether this a unique minimum or there are several quasi-stable states.

## Availability of data and materials

Data used in the preparation of this work were obtained from the OpenPain Project (OPP) database (www.openpain.org). The OPP project (Principal Investigator: A. Vania Apkarian, Ph.D. at Northwestern University) is supported by the National Institute of Neurological Disorders and Stroke (NINDS) and National Institute of Drug Abuse (NIDA). The preprocessing codes are available on request from the authors.

## Acknowledgements

We thank ACENET/Compute Canada for computational resources.

## Author Contributions

The planning of study, simulations and data analyses were done by GSM. GSM and JAH contributed equally in interpreting the results and writing of the paper.

## References

1. Loeser JD. Relieving pain in America [Internet]. Vol. 28, Clinical Journal of Pain. Washington, D.C.: National Academies Press; 2012 [cited 2019 Jan 22]. 185–186 p. Available from: http://www.nap.edu/catalog/13172

2. Hashmi JA, Baria AT, Baliki MN, Huang L, Schnitzer TJ, Apkarian VA. Brain networks predicting placebo analgesia in a clinical trial for chronic back pain. Pain [Internet]. 2012 Dec [cited 2019 Jan 22];153(12):2393–402. Available from: http://www.ncbi.nlm.nih.gov/pubmed/22985900

3. Hashmi JA, Baliki MN, Huang L, Baria AT, Torbey S, Hermann KM, et al. Shape shifting pain: Chronification of back pain shifts brain representation from nociceptive to emotional circuits. Brain [Internet]. 2013 [cited 2019 Jan 22];136(9):2751–68. Available from: www.iom.edu

4. Baliki MN, Chialvo DR, Geha PY, Levy RM, Harden RN, Parrish TB, et al. Chronic Pain and the Emotional Brain: Specific Brain Activity Associated with Spontaneous Fluctuations of Intensity of Chronic Back Pain. J Neurosci [Internet]. 2006 Nov 22 [cited 2019 Jan 22];26(47):12165–73. Available from: http://www.jneurosci.org/cgi/doi/10.1523/JNEUROSCI.3576-06.2006

5. Seminowicz DA, Wideman TH, Naso L, Hatami-Khoroushahi Z, Fallatah S, Ware MA, et al. Effective Treatment of Chronic Low Back Pain in Humans Reverses Abnormal Brain Anatomy and Function. J Neurosci [Internet]. 2011 May 18 [cited 2019 Jan 22];31(20):7540–50. Available from: http://www.ncbi.nlm.nih.gov/pubmed/21593339

6. Baliki MN, Geha PY, Apkarian A V., Chialvo DR. Beyond Feeling: Chronic Pain Hurts the Brain, Disrupting the Default-Mode Network Dynamics. J Neurosci [Internet]. 2008 Feb 6 [cited 2019 Jan 22];28(6):1398–403. Available from: http://www.ncbi.nlm.nih.gov/pubmed/18256259

7. Baliki MN, Schnitzer TJ, Bauer WR, Apkarian AV. Brain Morphological Signatures for Chronic Pain. Luque RM, editor. PLoS One [Internet]. 2011 Oct 13 [cited 2019 Jan 22];6(10):e26010. Available from: https://dx.plos.org/10.1371/journal.pone.0026010

8. Tagliazucchi E, Balenzuela P, Fraiman D, Chialvo DR. Brain resting state is disrupted in chronic back pain patients. Neurosci Lett [Internet]. 2010 [cited 2019 Jan 22];485(1):26–31. Available from: http://www.fmrib.ox.ac.uk/fsl

9. Yu R, Gollub RL, Spaeth R, Napadow V, Wasan A, Kong J. Disrupted functional connectivity of the periaqueductal gray in chronic low back pain. NeuroImage Clin [Internet]. 2014 [cited 2019 Jan 22];6:100–8. Available from: http://www.ncbi.nlm.nih.gov/pubmed/25379421

10. Tononi G, Edelman GM, Sporns O. Complexity and coherency: integrating information in the brain. Trends Cogn Sci [Internet]. 1998 Dec 1 [cited 2018 Oct 19];2(12):474–84. Available from: http://www.ncbi.nlm.nih.gov/pubmed/21227298

11. Crick F, Koch C. A framework for consciousness. Nat Neurosci [Internet]. 2003 Feb 1 [cited 2018 Oct 19];6(2):119–26. Available from: http://www.nature.com/articles/nn0203-119

12. Watts DJ, Strogatz SH. Collective dynamics of ‘small-world’ networks. Nature [Internet]. 1998 Jun 4 [cited 2019 Jan 22];393(6684):440–2. Available from: http://www.nature.com/articles/30918

13. Brody TA, Flores J, French JB, Mello PA, Pandey A, Wong SSM. Random-matrix physics: spectrum and strength fluctuations. Rev Mod Phys [Internet]. 1981 Jul 1 [cited 2019 Jan 22];53(3):385–479. Available from: https://link.aps.org/doi/10.1103/RevModPhys.53.385

14. Guhr T, Muïler-Groeling A, Weidenmuïler HA. Random-matrix theories in quantum physics: common concepts [Internet]. Vol. 299, Physics Reports. 1998 [cited 2019 Jan 22]. Available from: https://ac.els-cdn.com/S0370157397000884/1-s2.0-S0370157397000884-main.pdf?_tid=c74e85ab-91ea-4241-9c2c-bcbde9897d72&acdnat=1548187209_7eb224f592156ade5d7e052f74d2774b

15. Beenakker CWJ. Random-matrix theory of quantum transport. Rev Mod Phys [Internet]. 1997 Jul 1 [cited 2019 Jan 22];69(3):731–808. Available from: https://link.aps.org/doi/10.1103/RevModPhys.69.731

16. Bohigas O, Giannoni MJ, Schmit C. Characterization of Chaotic Quantum Spectra and Universality of Level Fluctuation Laws. Phys Rev Lett [Internet]. 1984 Jan 2 [cited 2019 Jan 22];52(1):1–4. Available from: https://link.aps.org/doi/10.1103/PhysRevLett.52.1

17. Seligman TH, Verbaarschot JJM, Zirnbauer MR. Quantum Spectra and Transition from Regular to Chaotic Classical Motion. Phys Rev Lett [Internet]. 1984 Jul 16 [cited 2019 Jan 22];53(3):215–7. Available from: https://link.aps.org/doi/10.1103/PhysRevLett.53.215

18. Bohigas O, Haq RU, Pandey A. Higher-Order Correlations in Spectra of Complex Systems. Phys Rev Lett [Internet]. 1985 Apr 15 [cited 2019 Jan 22];54(15):1645–8. Available from: https://link.aps.org/doi/10.1103/PhysRevLett.54.1645

19. Wintgen D, Marxer H. Level statistics of a quantized cantori system. Phys Rev Lett [Internet]. 1988 Mar 14 [cited 2019 Jan 22];60(11):971–4. Available from: https://link.aps.org/doi/10.1103/PhysRevLett.60.971

20. Pandey A, Ghosh S. Skew-Orthogonal Polynomials and Universality of Energy-Level Correlations. Phys Rev Lett [Internet]. 2001 Jun 21 [cited 2019 Jan 22];87(2):024102. Available from: https://link.aps.org/doi/10.1103/PhysRevLett.87.024102

21. Mehta ML. Random Matrices, Volume 142, Third Edition. Academic Press. 2004.

22. Santhanam MS, Patra PK. Statistics of atmospheric correlations. Phys Rev E [Internet]. 2001 Jun 11 [cited 2019 Jan 22];64(1):016102. Available from: https://link.aps.org/doi/10.1103/PhysRevE.64.016102

23. Plerou V, Gopikrishnan P, Rosenow B, Amaral LAN, Guhr T, Stanley HE, et al. Random matrix approach to cross correlations in financial data. Phys Rev E [Internet]. 2002 Jun 27 [cited 2018 Oct 19];65(6):066126. Available from: https://journals.aps.org/pre/pdf/10.1103/PhysRevE.65.066126

24. Sastry S, Deo N, Franz S. Spectral statistics of instantaneous normal modes in liquids and random matrices. Phys Rev E [Internet]. 2001 Jun 15 [cited 2019 Jan 22];64(1):016305. Available from: https://link.aps.org/doi/10.1103/PhysRevE.64.016305

25. Matharoo GS, Razul MSG, Poole PH. Spectral statistics of the quenched normal modes of a network-forming molecular liquid. J Chem Phys [Internet]. 2009 Mar 28 [cited 2018 Oct 19];130(12):124512. Available from: http://aip.scitation.org/doi/10.1063/1.3099605

26. Sarkar SK, Matharoo GS, Pandey A. Universality in the Vibrational Spectra of Single-Component Amorphous Clusters. Phys Rev Lett [Internet]. 2004 May 27 [cited 2018 Oct 19];92(21):215503. Available from: https://link.aps.org/doi/10.1103/PhysRevLett.92.215503

27. Matharoo GS, Sarkar SK, Pandey A. Vibrational spectra of amorphous clusters: Universal aspects. Phys Rev B [Internet]. 2005 Aug 1 [cited 2019 Jan 22];72(7):075401. Available from: https://link.aps.org/doi/10.1103/PhysRevB.72.075401

28. Matharoo GS. Universality in the Vibrational Spectra of Amorphous Systems. 2005 Dec 25 [cited 2019 Jan 22];(September). Available from: http://arxiv.org/abs/0812.4613

29. Matharoo GS. Universality in the vibrational spectra of weakly-disordered two-dimensional clusters. J Phys Condens Matter [Internet]. 2009 Feb 4 [cited 2018 Oct 19];21(5):055402. Available from: http://stacks.iop.org/0953-8984/21/i=5/a=055402?key=crossref.3ced8acd7b94f0348a398e2048dd0dfc

30. Osorio I, Lai Y-C. A phase-synchronization and random-matrix based approach to multichannel time-series analysis with application to epilepsy. Chaos An Interdiscip J Nonlinear Sci [Internet]. 2011 Sep [cited 2019 Jan 22];21(3):033108. Available from: http://aip.scitation.org/doi/10.1063/1.3615642

31. Bhadola P, Deo N. Targeting functional motifs of a protein family. Phys Rev E [Internet]. 2016 Oct 12 [cited 2019 Jan 22];94(4):1–13. Available from: https://link.aps.org/doi/10.1103/PhysRevE.94.042409

32. Šeba P. Random matrix analysis of human eeg data. Phys Rev Lett [Internet]. 2003 Nov 7 [cited 2019 Jan 22];91(19):1–4. Available from: https://link.aps.org/doi/10.1103/PhysRevLett.91.198104

33. Wang R, Zhang Z-Z, Ma J, Yang Y, Lin P, Wu Y. Spectral properties of the temporal evolution of brain network structure. Chaos An Interdiscip J Nonlinear Sci [Internet]. 2015 Dec 14 [cited 2018 Oct 19];25(12):123112. Available from: http://aip.scitation.org/doi/10.1063/1.4937451

34. Wang R, Wang L, Yang Y, Li J, Wu Y, Lin P. Random matrix theory for analyzing the brain functional network in attention deficit hyperactivity disorder. Phys Rev E [Internet]. 2016 Nov 23 [cited 2019 Jan 22];94(5):20–3. Available from: https://link.aps.org/doi/10.1103/PhysRevE.94.052411

35. Apkarian AV, Krauss BR, Fredrickson BE, Szeverenyi NM. Imaging the pain of low back pain: functional magnetic resonance imaging in combination with monitoring subjective pain perception allows the study of clinical pain states. Neurosci Lett [Internet]. 2001 Feb 16 [cited 2019 Jan 22];299(1–2):57–60. Available from: https://www.sciencedirect.com/science/article/pii/S030439400101504X

36. Biswal BB, Mennes M, Zuo X-N, Gohel S, Kelly C, Smith SM, et al. Toward discovery science of human brain function. Proc Natl Acad Sci [Internet]. 2010 [cited 2019 Jan 22];107(10):4734–9. Available from: https://www.pnas.org/content/pnas/107/10/4734.full.pdf

37. Hashmi JA, Loggia ML, Khan S, Gao L, Kim J, Napadow V, et al. Dexmedetomidine Disrupts the Local and Global Efficiencies of Large-scale Brain Networks. Anesthesiology [Internet]. 2017 [cited 2019 Jan 22];126(3):419–30. Available from: https://www.ncbi.nlm.nih.gov/pmc/articles/PMC5309134/pdf/nihms-835706.pdf

38. Hashmi JA, Kong J, Spaeth R, Khan S, Kaptchuk TJ, Gollub RL. Functional Network Architecture Predicts Psychologically Mediated Analgesia Related to Treatment in Chronic Knee Pain Patients. J Neurosci [Internet]. 2014 [cited 2019 Jan 22];34(11):3924–36. Available from: http://www.jneurosci.org/cgi/doi/10.1523/JNEUROSCI.3155-13.2014

39. Kelly C, Toro R, Di Martino A, Cox CL, Bellec P, Castellanos FX, et al. A convergent functional architecture of the insula emerges across imaging modalities. Neuroimage [Internet]. 2012 Jul 16 [cited 2019 Jan 24];61(4):1129–42. Available from: http://www.ncbi.nlm.nih.gov/pubmed/22440648

40. Rubinov M, Sporns O. Complex network measures of brain connectivity: Uses and interpretations. Neuroimage [Internet]. 2010 Sep [cited 2019 Jan 24];52(3):1059–69. Available from: http://www.ncbi.nlm.nih.gov/pubmed/19819337

41. Eguíluz VM, Chialvo DR, Cecchi GA, Baliki M, Apkarian AV. Scale-free brain functional networks. Phys Rev Lett. 2005;94(94):1–4.

42. Fraiman D, Balenzuela P, Foss J, Chialvo DR. Ising-like dynamics in large-scale functional brain networks. Phys Rev E - Stat Nonlinear, Soft Matter Phys. 2009;79(79):1–10.

43. Masuda N, Kojaku S, Sano Y. Configuration model for correlation matrices preserving the node strength. Phys Rev E [Internet]. 2018 Jul 20 [cited 2019 Jan 24];98(1):1–18. Available from: https://link.aps.org/doi/10.1103/PhysRevE.98.012312

44. Apkarian AV, Sosa Y, Sonty S, Levy RM, Harden RN, Parrish TB, et al. Chronic back pain is associated with decreased prefrontal and thalamic gray matter density. J Neurosci [Internet]. 2004 Nov 17 [cited 2019 Jan 22];24(46):10410–5. Available from: http://www.ncbi.nlm.nih.gov/pubmed/15548656

45. Cauda F, Costa T, Diano M, Sacco K, Duca S, Geminiani G, et al. Massive Modulation of Brain Areas After Mechanical Pain Stimulation: A Time-Resolved fMRI Study. Cereb Cortex [Internet]. 2014 Nov 1 [cited 2019 Jan 29];24(11):2991–3005. Available from: https://academic.oup.com/cercor/article-lookup/doi/10.1093/cercor/bht153

46. Borsook D, Edwards R, Elman I, Becerra L, Levine J. Pain and analgesia: the value of salience circuits. Prog Neurobiol [Internet]. 2013 May [cited 2019 Jan 29];104:93–105. Available from: http://www.ncbi.nlm.nih.gov/pubmed/23499729

47. Geuter S, Losin EAR, Roy M, Atlas LY, Schmidt L, Krishnan A, et al. Multiple brain networks mediating stimulus-pain relationships in humans. [cited 2019 Jan 29]; Available from: http://dx.doi.org/10.1101/298927

